# Morroniside promotes skin wound re-epithelialization by facilitating epidermal stem cell proliferation through GLP-1R-mediated upregulation of β-catenin expression

**DOI:** 10.1101/2024.02.06.579106

**Authors:** Chenghao Yu, Siyuan Yu, Zuohua Liu, Lei Xu, Zhiqiang Zhang, Jiaming Wan, Pengxiang Ji, Ping Zhang, Yi Fu, Yingying Le, Ruixing Hou

## Abstract

Epidermal stem cells (EpSCs) play a vital role in skin wound healing through re-epithelialization. Identifying chemicals that can promote EpSC proliferation is helpful for treating skin wounds. This study investigates the effect of morroniside on cutaneous wound healing in mice and explores the underlying mechanisms. Application of 10-50 μg/mL of morroniside to the skin wound promotes wound healing in mice. In vitro studies demonstrate that morroniside stimulates the proliferation of mouse and human EpSCs in a time- and dose-dependent manner. Mechanistic studies reveal that morroniside promotes the proliferation of EpSCs by facilitating the cell cycle transition from the G1 to S phase. Morroniside increases the expression of β-catenin via the glucagon-like peptide-1 receptor (GLP-1R)-mediated PKA, PKA/PI3K/AKT and PKA/ERK signalling pathways, resulting in the increase of cyclin D1 and cyclin E1 expression, either directly or by upregulating c-Myc expression. This process ultimately leads to EpSC proliferation. Administration of morroniside to mouse skin wounds increases the phosphorylation of AKT and ERK, the expression of β-catenin, c-Myc, cyclin D1, and cyclin E1, as well as the proliferation of EpSCs, in periwound skin tissue, and accelerates wound re-epithelialization. These effects of morroniside are mediated by the GLP-1R. Overall, these results indicate that morroniside promotes skin wound healing by stimulating the proliferation of EpSCs via increasing β-catenin expression and subsequent upregulation of c-Myc, cyclin D1, and cyclin E1 expression through GLP-1R signalling pathways. Morroniside has clinical potential for treating skin wounds.

## Introduction

The skin wound healing process consists of four sequential and overlapping phases: homeostasis, inflammation, proliferation, and remodeling. A variety of cells are involved in this process, including neutrophils, macrophages, and lymphocytes in the inflammatory phase, and epidermal stem cells/keratinocytes, fibroblasts, and vascular endothelial cells in the proliferative phase [1,2]. After a skin injury, epidermal stem cells (EpSCs) situated in the basal layer of the epidermis surrounding the wound edge proliferate, migrate towards the wound area, and differentiate into keratinocytes to repair the epidermis [3]. Stem cells located in the outer root sheath of hair follicles along the wound edge also participate in wound re-epithelialization [4]. To accelerate skin wound healing, research has mainly focused on regulating neutrophils, macrophages, and vascular endothelial cells to modulate inflammation and promote angiogenesis at the wound site [5–8]. However, there are limited reports on regulating the proliferation and differentiation of EpSCs for cutaneous wound re-epithelialization [9].

Morroniside is an iridoid glycoside extracted from the sarcocarp of Cornus officinalis [10]. It has been reported to promote angiogenesis in animal models of ischemic focal cerebral infarction [11,12] and myocardial infarction [13], as well as increase the survival of neurons and oligodendrocytes following spinal cord injury [14]. However, its effect on skin wound healing is currently unknown. Previous studies have shown that morroniside promotes cell proliferation [15–18]. Additionally, morroniside has been reported to alleviate neuropathic pain by activating spinal glucagon-like peptide-1 receptors (GLP-1R) [19]. GLP-1R agonists have been reported to stimulate cell proliferation, including mesenchymal stem cells and pancreatic β-cells [20,21]. Additionally, they have been found to facilitate skin wound healing [22,23]. GLP-1R is expressed in the epidermal cells of mouse skin and human keratinocytes [23]. Thus, we hypothesized that morroniside has the potential to improve skin wound healing by promoting wound re-epithelialization through GLP-1R.

In the present study, the effect of morroniside on the re-epithelialization and healing of skin wounds in mice was investigated through topical application of the compound. The effect and mechanisms of morroniside on EpSC proliferation were studied in vitro. Furthermore, the effect of morroniside on the proliferation of EpSCs residing in the skin epidermis at the wound edge, along with the molecules mediating this effect, was analyzed.

## Materials and Methods

### Chemicals

Morroniside was purchased from NatureStandard (Shanghai, China). 10058-F4, XAV-939, LY294002, PD98059, and exendin (9-39) were purchased from Selleck Chemicals LLC (Houston, USA). H89 2HCl was purchased from MedChemExpress LLC (New Jersey, USA). Thiazolyl blue tetrazolium bromide (MTT) was obtained from (Beyotime Biotechnology, Shanghai, China). Y-27632 was obtained from STEMCELL Technologies (Vancouver, Canada).

### Experiments on skin wound healing in mice

C57BL/6 mice were obtained from Changzhou Cavens Laboratory Animal Ltd. (Changzhou, China) and maintained in a conventional animal facility. The experimental protocols were reviewed and approved by the Ethics Committee of Suzhou Ruihua Orthopedic Hospital, Suzhou, China. Male mice aged 6-8 weeks were anesthetized with sevoflurane. A circular full-thickness skin defect wound was made using a 1.2 cm diameter biopsy punch (Haiyan Flagship Store, Suzhou, China) as previously described [24]. To investigate the effect of morroniside on wound healing, morroniside at concentrations of 10, 50, or 100 μg/mL, or the same volume of control vehicle (phosphate-buffered saline, PBS) was applied to the wound daily. To study the involvement of GLP-1R and downstream molecules in the improvement of wound healing by morroniside, the wounds were treated daily with PBS, 50 μg/mL morroniside, 415 μg/mL exendin (9-39), or a combination of 50 μg/mL morroniside with 415 μg/mL exendin (9-39). The wound area was photographed every other day. The wound size was measured utilizing the ImageJ software (NIH Image, Bethesda, USA). At 4 and 8 days post-treatment, the mice were anesthetized and sacrificed. A 0.5 cm wide region of skin tissue encompassing the wound was excised for subsequent histological, immunohistochemical, and Western blot analyses.

### Histological and immunohistochemical analyses

Mouse skin tissues were fixed in 10% formalin, embedded in paraffin and sectioned at 5 μm. Sections were deparaffinized and stained with hematoxylin and eosin to examine wound re-epithelialization. Immunohistochemistry was performed to detect β1 integrin- and PCNA-expressing cells in the regenerated epidermis as previously described [24]. Briefly, following heat-mediated antigen retrieval in sodium citrate using a microwave, serial skin sections were incubated with 0.3% H_2_O_2_ for 15 min, blocked with 10% goat serum for 2 h, and stained with a rabbit monoclonal antibody against β1 integrin and a mouse monoclonal antibody against PCNA (Abcam, Cambridge, UK) overnight at 4℃, respectively. After washing with PBS, the sections were incubated with a HRP-polymer anti-rabbit IgG or anti-mouse IgG secondary antibody (MXB Biotechnologies, Fuzhou, China) at room temperature for 1 h. The sections were then stained with 3’,3’-diaminobenzidine to detect β1 integrin or stained with 3-amino-9-ethylcarbazole to detect PCNA. Finally, the sections underwent counterstaining using hematoxylin and were photographed under a microscope. The thickness and area of the regenerated epidermis in H&E-stained sections, as well as the positive signals detected in immunohistochemically stained sections, were analyzed using ImageJ software.

### Isolation, culture and identification of EpSCs

Skin tissues were obtained from anesthetized neonatal C57BL/6 mice or from patients who underwent phase two plastic surgery following anterolateral femoral flap transplantation for skin injury treatment. The protocols were approved by the Ethics Committee of Suzhou Ruihua Orthopedic Hospital, Suzhou, China. EpSCs were isolated from skin tissue as previously described [24,25]. Briefly, the skin tissue was digested using 0.25% dispase II from Sigma-Aldrich (St Louis, USA) at 4°C for 13 h to separate the epidermis. The minced epidermis was incubated at 37°C for 15 min in 0.05% trypsin and then filtered through a 75 μm strainer. The resulting cell suspension was centrifuged, and the cell pellet was resuspended in Keratinocyte Growth Medium-2 Bullet Kit (KGM2 medium) consisting of KGM-2 Basal Medium and KGM-2 SingleQuots Supplements (Lonza, Basel, Switzerland), and then seeded onto Petri dishes coated with collagen IV. After a 10 min incubation at 37°C, the non-adherent cells were removed, and the adhering cells (EpSCs) were cultured in KGM2 medium containing 10 μM Y-27632. Subsequent experiments were conducted using the first passage mouse EpSCs and the second passage human EpSCs, both of which were cultured in KGM-2 basal medium.

Flow cytometry and immunofluorescence staining were conducted to detect EpSC biomarkers expression in cultured EpSCs as previously described [24,25]. In the flow cytometry assay, cultured mouse or human EpSCs were harvested and suspended in FACS buffer (1% goat serum and 5% FBS in PBS) at 25°C for 1 h. The cells were then incubated with FITC-conjugated α6 integrin antibody and PE-labelled CD71 antibody (BD Biosciences, San Jose, USA) at room temperature for 0.5 h. IgG2a labeled with either PE or FITC were utilized as isotype controls. The cells were washed and resuspended in FACS buffer before being analyzed through flow cytometry (Beckman Coulter, Pasadena, USA). For immunofluorescence staining, EpSCs were fixed in 10% paraformaldehyde for 15 min, permeabilized in 0.5% Triton X-100 for 15 min, and then blocked at room temperature using 10% goat serum for 2 h. The cells were incubated with a mouse monoclonal antibody targeting CK19 or a rabbit polyclonal antibody against β1 integrin (Abcam) overnight at 4°C, washed with PBS, and incubated with FITC-conjugated goat anti-mouse IgG or CY3 conjugated goat anti-rabbit IgG (Boster Bio, Wuhan, China) for 1 h. The cell nuclei were stained with DAPI (Beyotime Biotechnology) after washing. Fluorescence images were captured via a confocal immunofluorescence microscope (Zeiss, Oberkochen, Germany).

### Cell proliferation assays

EpSCs were either exposed to different concentrations of morroniside for varying durations or treated with inhibitors, including 3 μM exendin (9-39) for GLP-1R, 15 μM H89 for PKA, 20 μM LY294002 for PI3K, 20 μM PD98059 for ERK, 8 μM XAV-939 for β-catenin, and 2 μM 10058-F4 for c-Myc for 0.5 h followed by incubation with 20 μM morroniside or the same volume of PBS for 48 h. The proliferation of EpSCs was determined by MTT assay as previously described [25]. In addition, the effect of morroniside on the proliferation of EpSC was determined using the BrdU Cell Proliferation Kit (Abcam) according to the manufacturer’s instructions. The Multiskan Spectrum Microplate Reader (Thermo Fisher, Massachusetts, USA) was utilized to determine the optical density (OD) at 490 nm and 450 nm for the MTT assay and BrdU incorporation assay, respectively.

### Cell cycle analysis

Cultured EpSCs were collected and immobilized in 75% ethanol at 4°C overnight, centrifuged to remove ethanol, resuspended in PI/RNase staining buffer (BD Biosciences), and kept in the dark for 40 min. The DNA content of the cells was measured using a flow cytometer (Beckman Coulter, Pasadena, USA).

### Reverse transcriptase-polymerase chain reaction (RT-PCR)

Total RNA was extracted from mouse skin tissue or mouse/human EpSCs using Trizol reagent (Invitrogen, Massachusetts, USA), and used as a template for cDNA synthesize using HiScript II reverse transcriptase (Vazyme Biotech, Nanjing, China). PCR was conducted with 35 cycles of 95°C for 15 seconds, 60°C for 15 seconds, and at 72°C for 45 seconds. The PCR products were identified through agarose gel electrophoresis, which was followed by GelRed staining. The primer sequences (5′-3′) of the PCR are as follows: human *GLP-1R*: 5′-CAGCGCTCCCTGACTGAG (forward), CAGGCGTATTCATCGAAGGT (reverse); mouse *Glp-1r*: TTTCCTCACGGAAGCGCCACTCC (forward), GGATAACGAACAGCAGCGGAACTCCC (reverse), human β-actin: GGCTGTGCTATCCCTGTACG (forward), CTTGATCTTCATTGTGCTGGGTG (reverse), mouse *Gapdh*: GGTCCCAGCTTAGGTTCATCA (forward), CCTTTTGGCTCCACCCTTCA (reverse).

### Western blot analysis

Proteins were extracted from skin tissues or cultured EpSCs with RIPA lysis buffer containing Protease and Phosphatase Inhibitor Cocktail (Beyotime Biotechnology). Protein concentrations were determined by using the BCA Protein Quantification Kit (Vazyme Biotech). Western blot was carried out following the standard protocol. Briefly, proteins separated by SDS-PAGE were transferred to a PVDF membrane. After blocking with 5% nonfat milk (Sangon Biotech, Shanghai, China), the membrane was incubated with primary antibodies targeting phospho-AKT, AKT, phospho-ERK1/2, phospho-β-catenin (Cell Signaling Technology, Massachusetts, USA), ERK1/2, β-catenin, c-Myc, cyclin E1, cyclin D1 (Abcam,), and GAPDH (Proteintech, Chicago, USA), washed and incubated with secondary antibodies. The High-Sensitivity ECL Chemiluminescence Detection Kit from Vazyme Biotech was used to detect target proteins, and quantification was performed through ImageJ software (NIH Image).

### Statistical analysis

Each experiment was replicated a minimum of three times. Data are presented as the mean ± standard deviation (SD) or standard error of the mean (SEM). The data were analyzed using GraphPad Prism 9 software. Statistical significance between two groups was determined using an unpaired Student’s t-test. A P value of 0.05 or less was considered statistically significant.

## Results

### Morroniside promotes cutaneous wound healing in mice

To investigate the effect of morroniside on skin wound healing in mice, different concentrations of morroniside were applied to the wound daily. Morroniside at 10 and 50 μg/mL was found to significantly promote skin wound healing (Figure 1A, B). After 12 days of treatment, H&E staining revealed that both 10 and 50 μg/mL of morroniside significantly increased the epidermal thickness of the regenerated skin (Figure 1C,D). These results suggest that morroniside promotes skin wound healing by promoting re-epithelialization of the wound.

**Figure 1.**
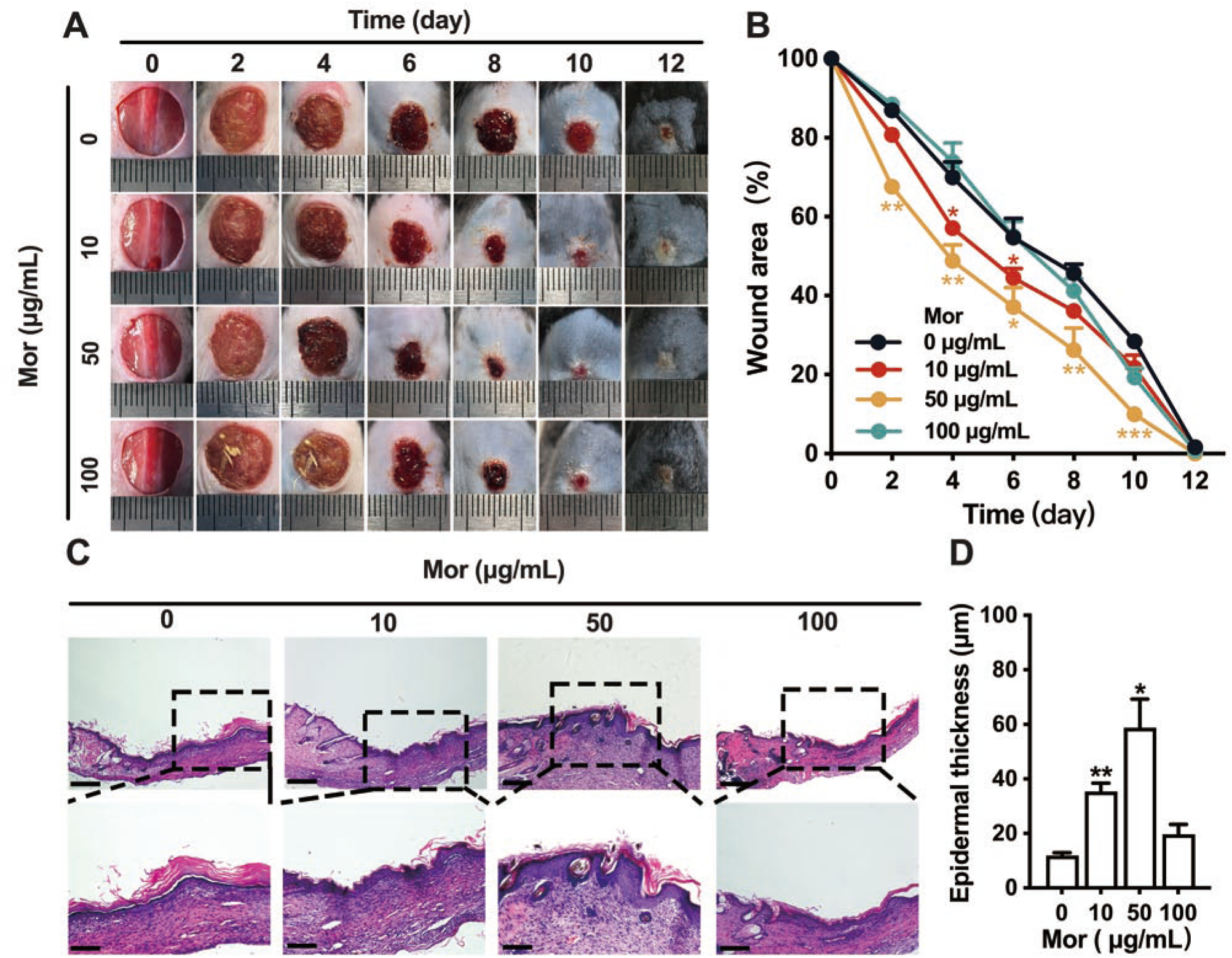
Morroniside promotes skin wound healing in mice. (A-B) Representative photographs of skin wound (A) and changes in wound area (B) after treatment with different concentrations of morroniside (Mor) or PBS. (C-D) Representative H&E staining images of regenerated skin tissue (C) and the thickness of regenerated epidermis at day 12 after treatment with different concentrations of Mor or PBS. n=4 mice/group, data are mean ± SE. *p < 0.05, **p < 0.01, ***p < 0.001, compared with PBS-treated mice. Scale bars: 100 μm.

### Morroniside enhances the proliferation of EpSC by promoting cell cycle progression from G1 to S phase

Since EpSCs play a crucial role in wound re-epithelialization, we examined the effect of morroniside on the proliferation of EpSCs in vitro. Cultured EpSCs, isolated from either mouse or human skin epidermis, display a cobblestone-like appearance under light microscopy (Figure 2A,D). These cells exhibited high α6 integrin expression and low CD71 levels (Figure 2B,E), and were positive for β1 integrin and cytokeratin 19 (CK19) (Figure 2C,F), which are biomarkers of EpSCs. MTT assay showed that treatment of mouse EpSCs with 5-40 μM morroniside significantly increased cell proliferation after 48 h (Figure 3A). The maximum proproliferative effect was observed with 20 μM morroniside, and this concentration exhibited a time-dependent stimulation of mouse EpSC proliferation (Figure 3A,B). The proproliferative effect of 20 μM morroniside on mouse EpSCs was validated by the BrdU incorporation assay (Figure. 3C). MTT assay (Figure 3D,E) and BrdU incorporation assays (Figure 3F) showed that morroniside also induced human EpSC proliferation in a dose- and time-dependent manner.

**Figure 2.**
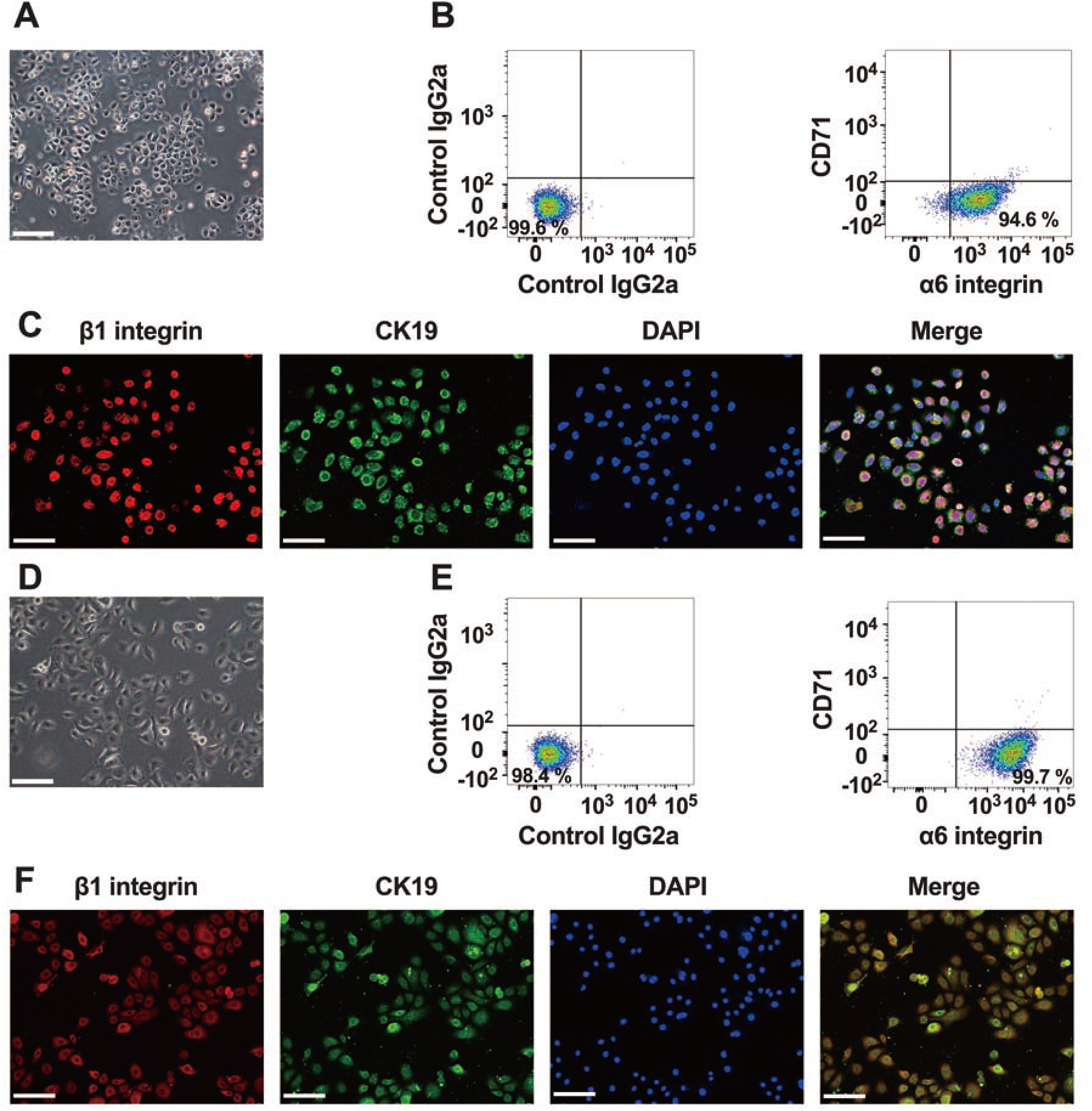
Characterization of mouse and human EpSCs. (A, D) Morphology of mouse EpSCs (A) and human EpSCs (D) under a light microscope. (B, E) Flow cytometry analysis of α6 integrin and CD71 expression in mouse EpSCs (B) and human EpSCs (E). (C, F) Immunofluorescence staining of CK19 (green) and β1 integrin (red) in mouse EpSCs (C) and human EpSCs (F). Cell nuclei were stained with DAPI. Scale bars: 100 μm. The photographs of mouse and human EpSCs, immunofluorescence staining images, and flow cytometry results are representative of three independent experiments.

**Figure 3.**
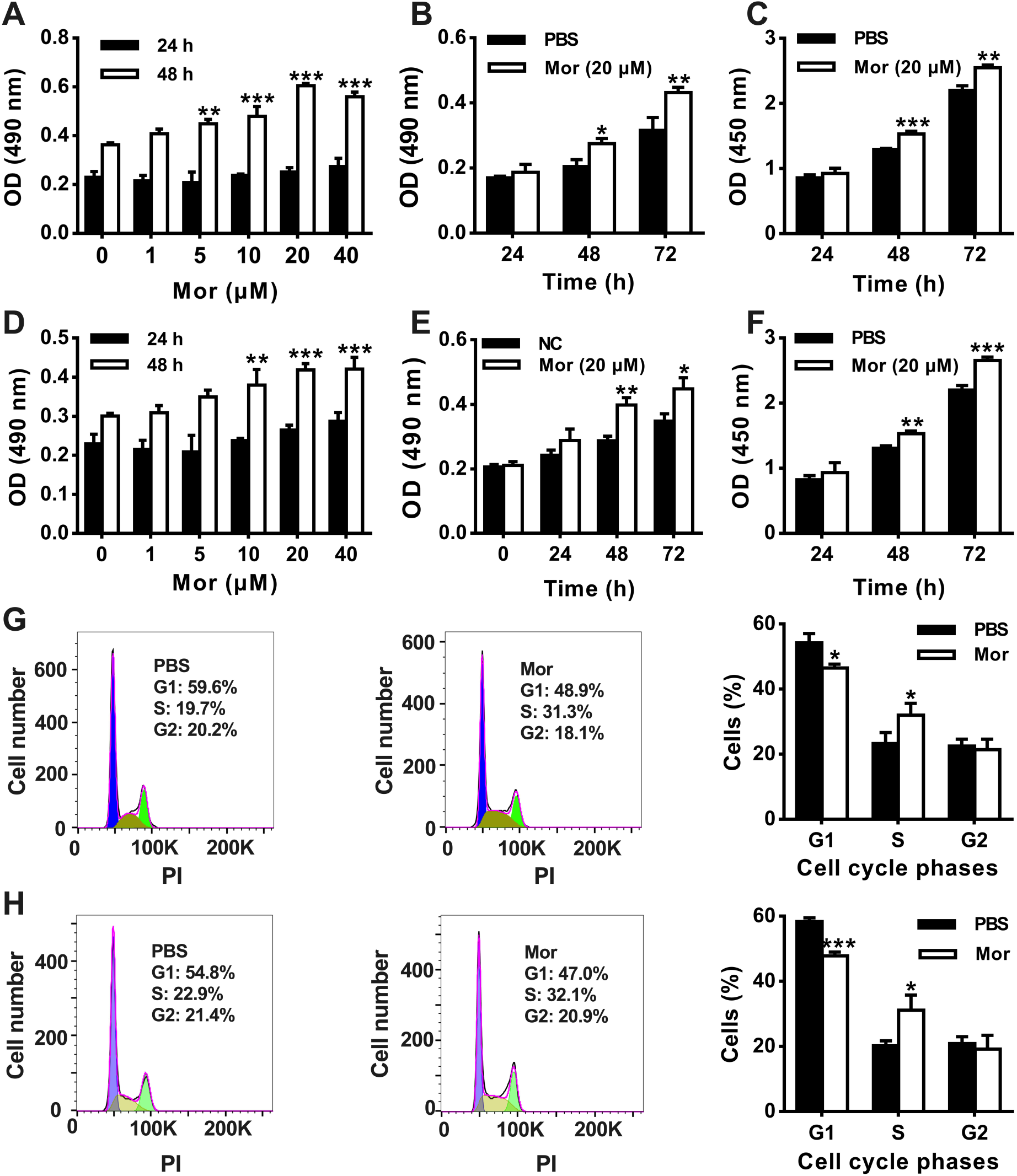
Morroniside induces EpSC proliferation by promoting the G1/S cell cycle transition. (A-F) Proliferation of mouse EpSCs (A-C) and human EpSCs (D-F) treated with different concentrations of morroniside (Mor) or PBS for 24 h and 48 h (A, D) was determined by MTT assay, treated with 20 μM morroniside or equal volumes of PBS for different time periods was determined by MTT assay (B, E) and BrdU incorporation assay (C, F), respectively. (G-H) Cell cycle distribution of mouse EpSCs (G) and human EpSCs (H) treated with 20 μM morroniside or the same volume of PBS for 48 h was determined by flow cytometry. n=3, data are mean ± SD. *p < 0.05, **p < 0.01, ***p < 0.001, compared with cells treated with PBS for the same period of time.

We then examined the effect of morroniside on the cell cycle of EpSCs. Analysis using flow cytometry revealed that morroniside treatment led to a significant decrease in the proportion of cells in the G1 phase and a remarkable increase in the proportion of cells in the S phase in EpSCs from both mice and humans (Figure 3G,H), indicating that morroniside promotes the proliferation of EpSCs by facilitating the cell cycle transition from the G1 phase to the S phase.

### Morroniside promotes EpSC proliferation by activating the β-catenin and β-catenin/c-Myc signaling pathways through GLP-1R

Since β-catenin and c-Myc play significant roles in EpSC proliferation [26,27], and cyclin D1 and cyclin E1 contribute to the cell cycle transition from G1 phase to S phase [28,29], we investigated the effect of morroniside on the expression of these molecules in mouse EpSCs. Western blot assay showed that 20 μM morroniside upregulated the expression of β-catenin, c-Myc, cyclin Dl, and cyclin El in a time-dependent manner (Figure 4A). The MTT assay showed that pretreating mouse EpSCs with either β-catenin signalling pathway inhibitor XAV-939 or the c-Myc inhibitor 10058-F4 significantly suppressed cell proliferation induced by morroniside (Figure 4B,D). Western blot assay showed that the upregulation of β-catenin, c-Myc, cyclin Dl, and cyclin El induced by morroniside was significantly inhibited by XAV-939 (Figure 4C). Additionally, morroniside-induced expression of c-Myc, cyclin Dl, and cyclin El was strongly inhibited by 10058-F4 (Figure 4E). These findings indicate that morroniside enhances the proliferation of EpSCs by increasing the expression of cyclin D1 and cyclin E1 through the activation of β-catenin and β-catenin/c-Myc signalling pathways.

**Figure 4.**
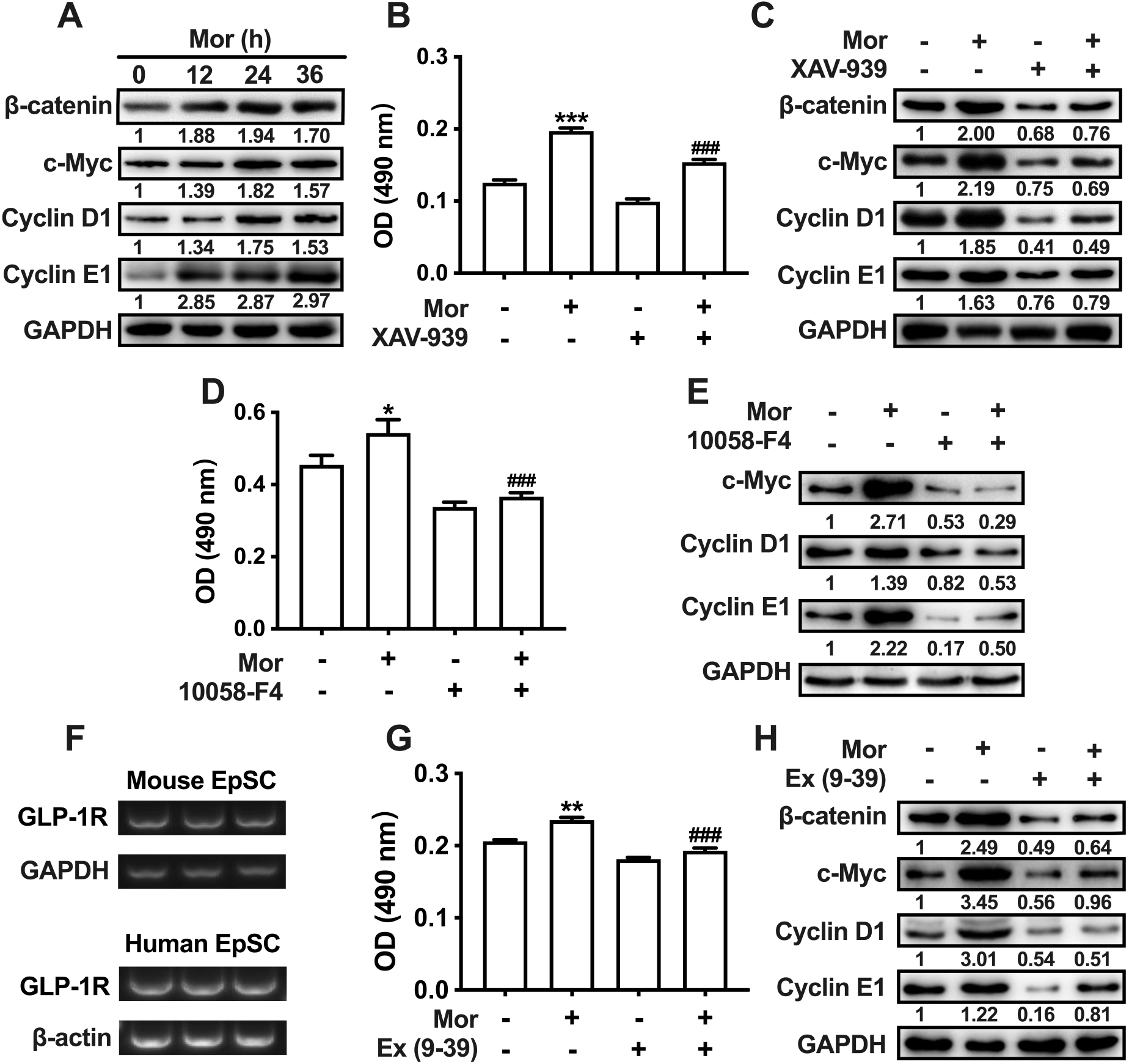
Morroniside promotes EpSC proliferation through GLP-1R-mediated upregulation of β-catenin, c-Myc and cyclins. (A) Western blot detection of β-catenin, c-Myc, cyclin Dl, and cyclin E1 expression in mouse EpSCs treated with 20 μM morroniside (Mor) for different time periods. (B-E,G,H) Mouse EpSCs pretreated with/without 8 μM XAV-939, 2 μM 10058-F4, or 3 μM exendin (9-39) (Ex (9-39)) for 0.5 h were incubated with 20 μM monosodium or same volume of PBS and examined for cell proliferation after 48 h by MTT assay (B, D, G), for expression of β-catenin, c-Myc, cyclin Dl, and cyclin E1 after 24 h by Western blot (C, E, H). (F) Detection of GLP-1R mRNA expression in mouse and human EpSCs by RT-PCR. n=3, data are mean ± SD. *p < 0.05, **p < 0.01, ***p < 0.001, compared with PBS-treated cells. ^###^p < 0.001, compared with cells treated with Mor. A, C, E and H are representative results from three replicate experiments.

It has been reported that morroniside is a GLP-1R agonist [19]. We investigated whether morroniside stimulates EpSC proliferation through GLP-1R. RT-PCR assay showed that both mouse and human EpSCs expressed GLP-1R (Figure 4F). MTT assay and Western blot assay showed that pretreatment of mouse EpSCs with the GLP-1R antagonist exendin (9-39) (Ex (9-39)) significantly inhibited cell proliferation (Figure 4G), as well as expression of β-catenin, c-Myc, cyclin Dl, and cyclin El induced by morroniside (Figure 4H). This suggests that morroniside promotes EpSC proliferation by upregulating the expression of β-catenin, c-Myc, cyclin Dl, and cyclin El via GLP-1R. Taken together, the above-mentioned findings indicate that morroniside enhances EpSC proliferation by increasing the expression of cyclin D1 and E1 through GLP-1R-mediated β-catenin and β-catenin/c-Myc pathways.

### Morroniside promotes EpSC proliferation through GLP-1R/PKA, GLP-1R/PKA/PI3K/AKT and GLP-1R/PKA/ERK signaling pathways mediated upregulation of β-catenin

GLP-1R agonist GLP-1 has been reported to stimulate pancreatic β-cell proliferation through protein kinase A (PKA), PI3K, and ERK [21]. As morroniside functions as an agonist of GLP-1R [19], we investigated if these molecules also contribute to EpSC proliferation induced by morroniside. The MTT and Western blot assays showed that PKA inhibitor H89, PI3K inhibitor LY294002, or ERK inhibitor PD98059 suppressed the proliferation of mouse EpSCs (Figure 5A-C) and the expression of β-catenin, c-Myc, cyclin Dl, and cyclin El expression induced by morroniside (Figure 5D). These results indicate that morroniside induces EpSCs proliferation by increasing the expression of β-catenin, c-Myc, cyclin Dl, and cyclin El through PKA, PI3K and ERK signalling pathways. A time course study showed that morroniside significantly increased AKT and ERK phosphorylation in mouse EpSCs after 24 h of stimulation (Figure 5E). Pretreatment of mouse EpSCs with Ex (9-39) or H89 suppressed morroniside-induced AKT and ERK phosphorylation (Figure 5F,G). This indicates that morroniside activates AKT and ERK through GLP-1R and PKA. Collectively, the above findings indicate that morroniside enhances EpSC proliferation by increasing the expression of β-catenin, c-Myc, cyclin D1 and E1 through the GLP-1R/PKA, GLP-1R/PKA/PI3K/AKT and GLP-1R/PKA/ERK signalling pathways.

**Figure 5.**
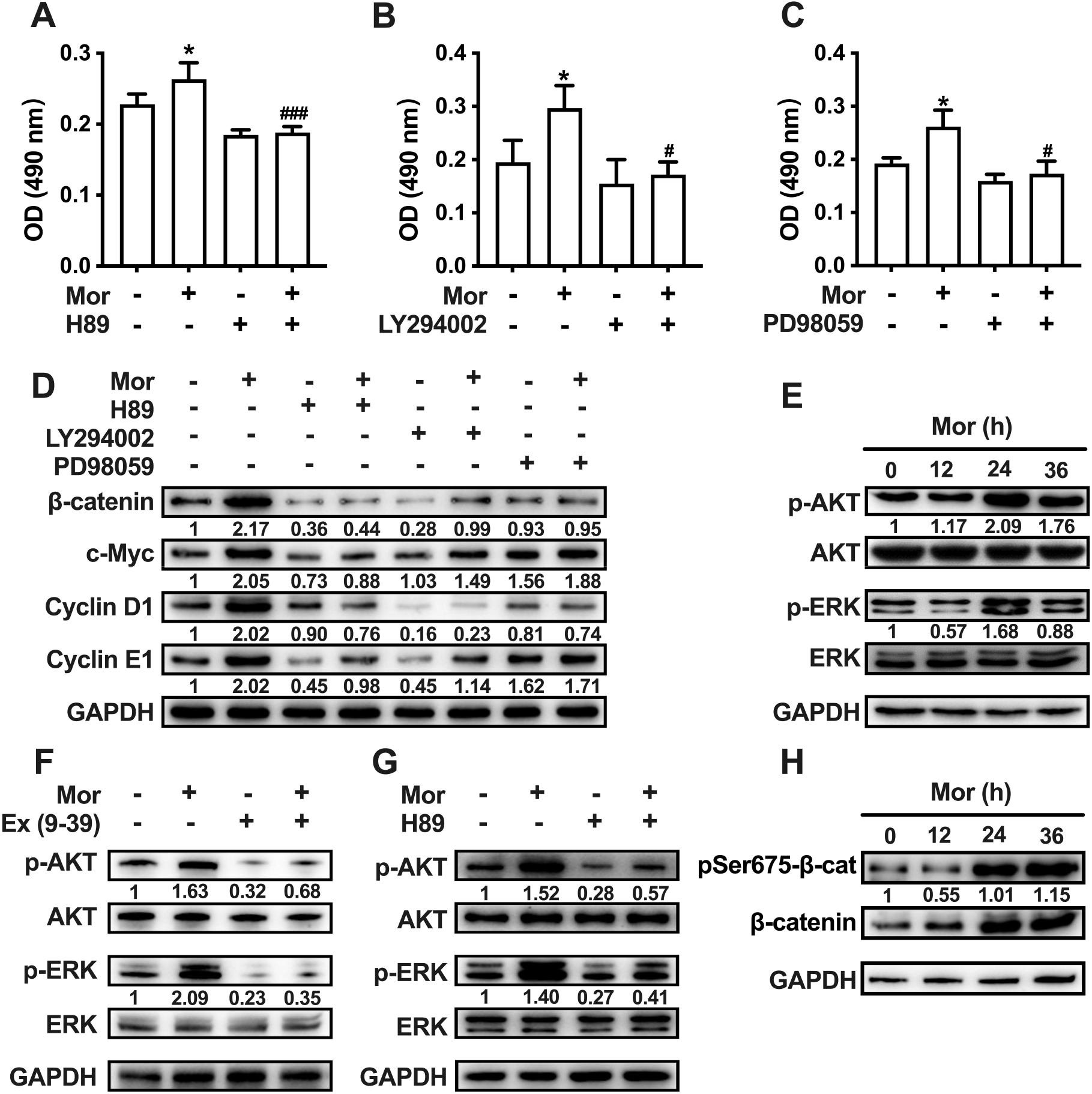
Morroniside stimulates EpSC proliferation by enhancing β-catenin expression through the GLP-1R/PKA, GLP-1R/ PKA/PI3K/AKT and GLP-1R/PKA/ERK pathways. (A-D,F,G) Mouse EpSCs pretreated with/without 15 μM H89, 20 μM LY294002, 20 μM PD98059, or 3 μM exendin (9-39) (Ex (9-39)) for 0.5 h were incubated with 20 μM morroniside (Mor) or an equal volume of PBS, cell proliferation was determined by MTT assay 48 h later (A-C), the expression of β-catenin, c-Myc, cyclin D1 and cyclin E1 (D), as well as the phosphorylation of AKT and ERK (F, G) were detected by Western blot 24 h later. (E,H) The phosphorylation of AKT, ERK (E) and β-catenin at Ser675 (H) in mouse EpSCs treated with 20 μM morroniside for different time periods was analyzed by Western blot. A-C: n=3, data are mean ± SD. *p< 0.05, compared with PBS-treated cells. ^#^p < 0.05, ^###^p < 0.001, compared with Mor-treated cells. D-J are representative results of three replicate experiments.

As PKA activation has been reported to increase the transcriptional activity of β-catenin by phosphorylating it at serine 675 (Ser675) and stabilizing the protein [30,31]. We investigated whether morroniside could promote Ser675 phosphorylation on β-catenin in mouse EpSCs through the GLP-1R/PKA signaling pathway. A time course study showed that morroniside increased both β-catenin expression and phosphorylation at Ser675 after a 24 h treatment (Figure 5H). However, morroniside treatment did not significantly increase the ratio of phosphorylated β-catenin to total β-catenin (Figure 5H), indicating that morroniside has no significant effect on β-catenin phosphorylation.

### Morroniside enhances EpSC proliferation and skin wound re-epithelialization via GLP-1R and downstream molecules in mice

To investigate whether morroniside stimulates EpSC proliferation via the GLP-1R signaling pathway to promote epidermal regeneration of skin wounds, mouse skin wounds were treated daily with PBS, morroniside, the GLP-1R antagonist Ex (9-39), or a combination of morroniside with Ex (9-39). The wound area was photographed every other day, and the skin tissues surrounding the wound were collected on day 4 and day 8 after treatment. The Ex (9-39) had no significant effect on wound healing; however, it reversed the accelerating effect of 50 μg/ml morroniside on wound healing (Figure 6A, B). H&E staining showed that morroniside greatly increased the thickness and size of the regenerated epidermis after 4 and 8 days of treatment. The application of Ex (9-39) notably suppressed the effect of morroniside (Figure 6C-E), implying that morroniside accelerates the re-epithelialization of wounds via GLP-1R. Immunohistochemical staining for the EpSC biomarker β1 integrin and the cell proliferation marker proliferating cell nuclear antigen (PCNA) in consecutive skin tissue sections demonstrated that morroniside substantially increased the number of cells positive for β1 integrin or PCNA in the basal layer of regenerated epidermis. The majority of these cells were double-positive for β1 integrin and PCNA. Ex (9-39) markedly reduced the increase of β1 integrin and PCNA single-positive cells, as well as double-positive cells, induced by morroniside (Figure 7A,B). These findings suggest that morroniside stimulate EpSC proliferation through GLP-1R during wound healing. Western blot analysis showed that after 4 days of treatment with morroniside, there was a significant increase in the phosphorylation of AKT and ERK (Figure 7C,D) and the expression of β-catenin, c-Myc, cyclin D1, and cyclin E1 (Figure 7E,F) in skin tissues at the wound edge. Treatment with Ex (9-39) considerably reduced the positive effect of morroniside on either the expression or the phosphorylation of these molecules (Figure 7C-F). These results indicate that morroniside promotes the proliferation of EpSCs at the wound edge through GLP-1R-mediated activation of AKT and ERK, upregulation of β-catenin, c-Myc, cyclin D1, and cyclin E1 expression. Collectively, our findings demonstrate that morroniside accelerates re-epithelialization of skin wounds by stimulating EpSC proliferation through GLP-1R and downstream molecules.

**Figure 6.**
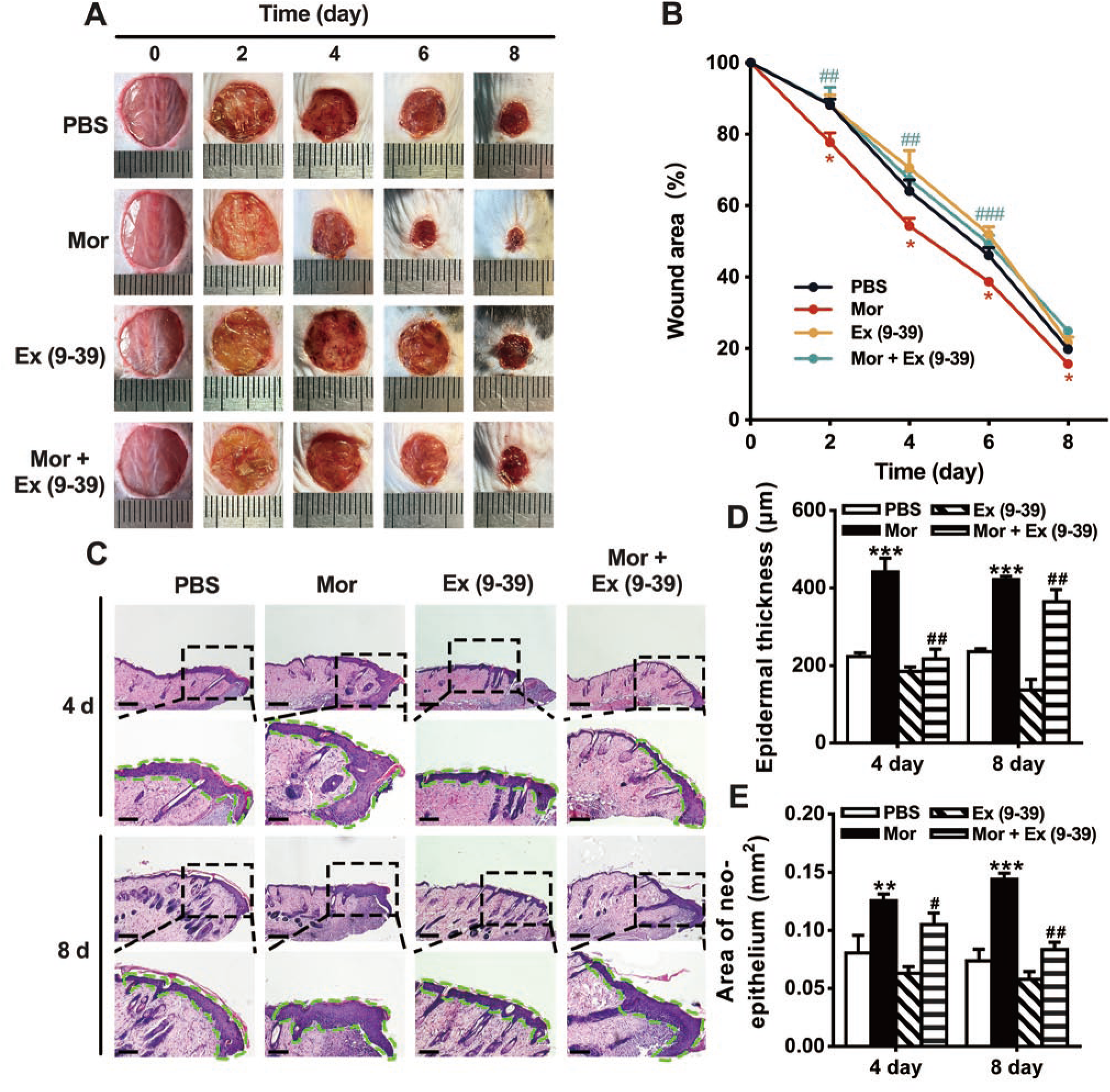
Morroniside promotes skin wound re-epithelialization via GLP-1R. (A-B) Representative images of skin wound (A) and changes in wound area (B) after treatment with PBS, 50 μg/mL morroniside (Mor), 415 μg/mL exendin (9-39) (Ex (9-39)), or 50 μg/mL Mor in combination with 415 μg/mL Ex (9-39). (C-E) Representative images of H&E staining of periwound skin tissue on days 4 and 8 after treatment (C) and quantitative analysis of thickness (D) and area of regenerated epidermis (E). n=4 mice/group, data are mean ± SE. *p < 0.05, **p < 0.01, ***p < 0.001, compared with PBS-treated mice. ^#^p < 0.05, ^##^p < 0.01, ^###^p < 0.001, compared with Mor-treated mice. Scale bars: 100 μm.

**Figure 7.**
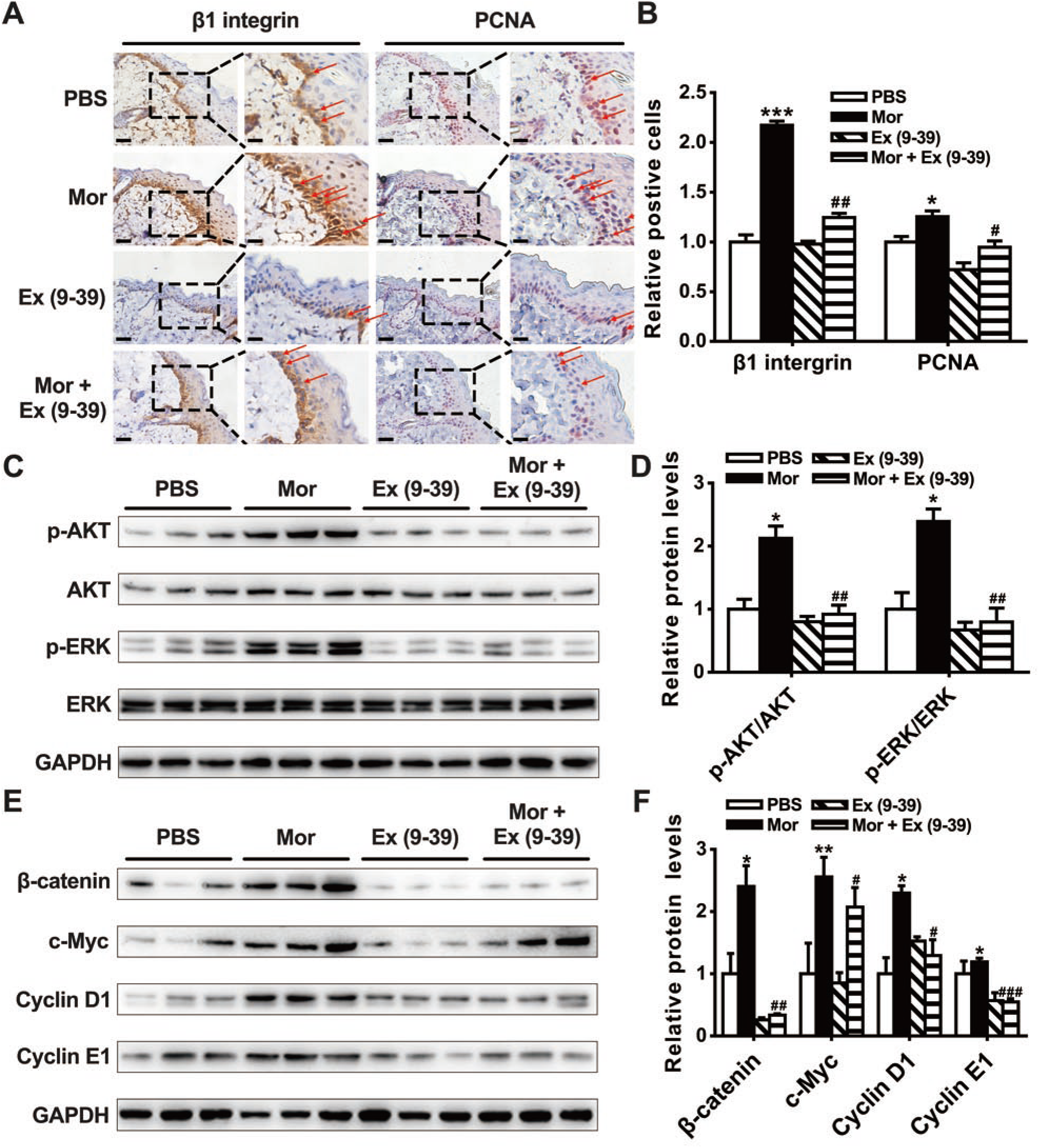
Morroniside promotes EpSC proliferation in mouse skin wound through GLP-1R-mediated activation of downstream molecules. Mouse skin wounds were treated with PBS, 50 μg/mL morroniside (Mor), 415 μg/mL exendin (9-39) (Ex (9-39)), or 50 μg/mL Mor together with 415 μg/mL Ex (9-39) for 4 days. Skin around the wound edge was collected and examined for EpSC proliferation by immunohistochemical staining for β1 integrin and PCNA-expressing cells in serial sections (A, B), and for phosphorylation of AKT and ERK (C, D), expression of β-catenin, c-Myc, cyclin D1, and cyclin E1 (E, F) by Western blot assay. Scale bars are 40 μm and 20 μm in the left and right panels of β1 integrin and PCNA immunohistochemistry images, respectively. Arrows indicate β1 integrin and PCNA double-positive cells. n=3 mice/group. Data are mean ± SE, *p < 0.05, **p < 0.01, ***p < 0.001, compared with PBS-treated mice; ^#^p < 0.05, ^##^p < 0.01, ^###^p < 0.001, compared with Mor-treated mice.

## 4. Discussion

In this study, we investigated the effect of morroniside on the healing of cutaneous wounds in mice and explored the underlying mechanisms. We provided evidence that morroniside promoted skin wound re-epithelialization and healing. Mechanistic studies demonstrated that morroniside promoted the proliferation of EpSCs by facilitating the cell cycle transition from the G1 to S phase. Morroniside enhanced EpSC proliferation through the GLP-1R-mediated activation of β-catenin signalling via the PKA, PKA/PI3K/AKT and PKA/ERK signalling pathways. An in vivo study confirmed that morroniside increased the proliferation of EpSCs and facilitated the re-epithelialization of skin wounds in mice via GLP-1R and downstream molecules.

Using a murine model of full-thickness skin defects, we found that the administration of morroniside to skin wounds accelerated wound healing by promoting wound re-epithelialization (Figure 1, Figure 6). In vitro studies using EpSCs isolated from both mouse and human skin tissue demonstrated that morroniside stimulated EpSC proliferation. Treatment of EpSCs with morroniside promoted cell cycle transition from the G1 phase to S phase (Figure 3G, H). It is well known that cell cycle progression is mainly regulated by cyclin-dependent kinases (CDKs) and cyclins. Cyclin D binds to CDK4 or CDK6 to promote cells to transition from quiescence into the cell cycle and progress through G1 phase. Cyclin E binds to CDK2 to control cell cycle progression from the G1 phase to the S phase [32]. Our study showed that morroniside increased the expression of cyclin D1 and cyclin E1 in mouse EpSCs (Figure 4A). This indicate that by enhancing the expression of cyclin D1 and cyclin E1, morroniside facilitates the transition of cell cycle from G1 phase to S phase, thereby promoting EpSC proliferation.

The Wnt/β-catenin signalling pathway plays a significant role in the self-renewal and proliferation of skin EpSCs [26,33,34]. Activation of this signaling pathway following skin injury promotes EpSC proliferation and wound re-epithelialization [35,36]. c-Myc, a downstream molecule of β-catenin [37], is a transcription factor that induces the expression of several positive regulators of the cell cycle, such as CDKs, cyclins, and E2F transcription factors [38]. Our data demonstrated that morroniside promotes the expression of β-catenin, c-Myc, cyclin D1, and cyclin E1 in EpSCs in a time-dependent way. XAV-939, a β-catenin signalling inhibitor that stimulates β-catenin degradation by stabilizing axin via inhibiting tankyrase 1 and tankyrase 2 [39], and 10058-F4, a c-Myc inhibitor, were used to examine the role of β-catenin and c-Myc, in morroniside-induced proliferation of EpSC and expression of cyclin D1 and cyclin E1 (Figure 4B-E). The results indicate that morroniside promotes the proliferation of EpSC by enhancing the expression of cyclin D1 and cyclin E1 through both β-catenin and β-catenin/c-Myc signalling pathways.

Morroniside is an orthosteric agonist of GLP-1R [19]. GLP-1R agonist has been shown to enhance pancreatic β-cells proliferation by upregulating cyclin D1 expression through PKA, PI3K and ERK pathways [21]. We observed that both mouse and human EpSCs expressed GLP-1R (Figure 4F). Experiments conducted with GLP-1R antagonist Ex (9-39), PKA inhibitor H89, PI3K inhibitor LY294002, and ERK inhibitor PD98059 demonstrated the inhibition of EpSC proliferation (Figure 4G, Figure 5A-C) and β-catenin, c-Myc, cyclin D1 and cyclin E1 expression induced by morroniside (Figure 4H, Figure 5D). Additionally, Ex (9-39) and H89 have been shown to inhibit morroniside-induced AKT and ERK phosphorylation in EpSCs (Figure 5F,G). These findings indicate that morroniside activates GLP-1R and downstream PKA, PKA/PI3K/AKT and PKA-ERK signaling pathways, leading to the expression of β-catenin, c-Myc, cyclin D1, and cyclin E1, which in turn promotes EpSC proliferation. Figure 5D shows that the PKA inhibitor had stronger inhibitory effect on the expression of β-catenin, c-Myc, and cyclin E1 induced by morroniside compared to the AKT inhibitor and ERK inhibitor. This result suggests that the GLP-1R/PKA signaling pathway may play a more significant role than the other two pathways in mediating the proproliferative effect of morroniside on EpSCs. Previous studies have reported that the activation of PI3K/AKT prevents the degradation of β-catenin, c-Myc and cyclin D by phosphorylating glycogen synthase kinase β and inactivating it [40,41]. ERK activation has been found to phosphorylate serine 62 of c-Myc, resulting in an increase in its stability [40]. In addition, ERK activation has been shown to regulate G1/S transition by increasing the transcription of cyclin D1 and cyclin E [42,43]. These mechanisms may be also be responsible for the increase in protein levels of β-catenin, c-Myc, cyclin D1, and cyclin E1 in EpSCs following treatment with morroniside but require further investigation.

Finally, our in vivo study demonstrated that morroniside enhanced skin wound re-epithelialization in mice. It stimulated the phosphorylation of AKT and ERK, increased the protein levels of β-catenin, c-Myc, cyclin D1 and cyclin E1, and the proliferation of EpSCs in the epidermis at the wound edge in mice via GLP-1R. These findings provide evidence for the effectiveness of morroniside in activating GLP-1R signaling pathways to upregulating the expression of β-catenin and downstream molecules, thereby stimulating EpSC proliferation and facilitating skin wound healing.

In summary, our study demonstrates that morroniside promotes skin wound healing in mice by stimulating the proliferation of EpSCs and the re-epithelialization of wounds. Morroniside increases β-catenin expression through GLP-1R-mediated PKA, PKA/PI3K/AKT, and PKA/ERK signalling pathways in EpSCs. This leads to an elevation in β-catenin transcriptional activity, which initiates the transcription of c-Myc, cyclin D1 and cyclin E1. c-Myc also induces the expression of cyclin D1 and cyclin E1. Consequently, the upregulation of c-Myc, along with cyclin D1 and cyclin E1, induces the proliferation of EpSCs (Figure 8). Morroniside has the potential to serve as a therapeutic agent for treating skin wounds.

**Figure 8.**
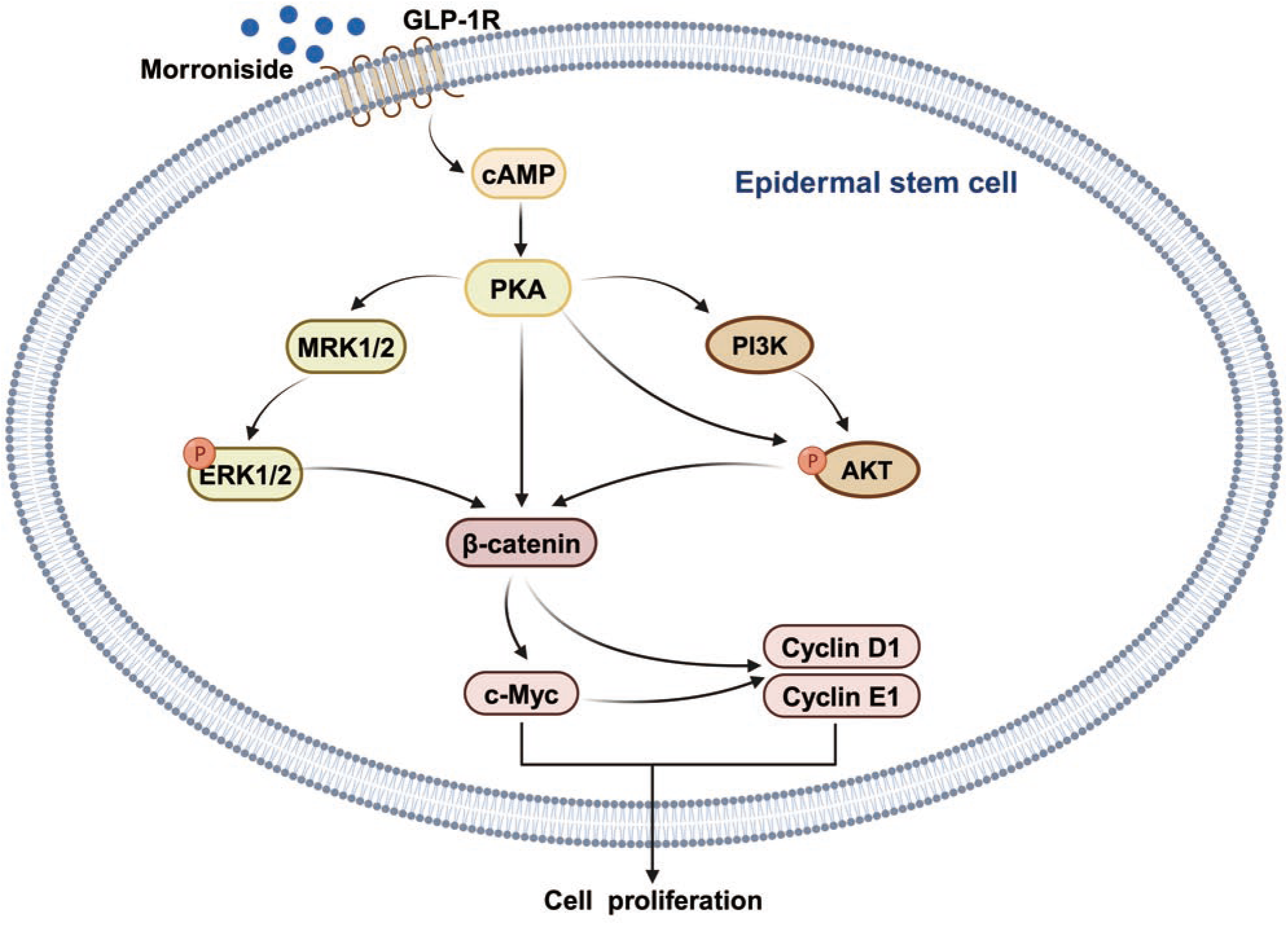
An illustration of the signalling pathways involved in the proliferation of EpSCs induced by morroniside. AKT, protein kinase B; ERK, extracellular signal-regulated kinase; EpSC, epidermal stem cell; GLP-1R, glucagon-like peptide 1 receptor; PI3K, phosphatidylinositol-4,5-bisphosphate 3-kinases; PKA, protein kinase A.

